# Role of tolerance to resource demand – supply mismatch in a model of annual plants

**DOI:** 10.1101/075804

**Authors:** Michel Droz, Andrzej Pękalski

**Affiliations:** Department of Theoretical Physics, University of Geneva, 1211 Genéve 4, Switzerland; Institute of Theoretical Physics, University of Wrocław, pl. M. Borna 9, 50-203 Wrocław, Poland

**Keywords:** Monte Carlo simulations, plant dynamics, annual plants, coexistence, resource shortage or surplus

## Abstract

We propose and discuss a simulation model of annual plants competing for a single resource. Plants are characterised by their tolerance to a shortage of it and the maximum number of seeds a plant could produce in a year. Interaction among plants is reduced to blocking a part of the resource by the plant’s nearest neighbours. Spatial and temporal external conditions remain unchanged. We consider two systems - plants of one type (target plants) and a mixture of two types, when the second type differs from the target ones only by the tolerance to a shortage or surplus of the resource. We show how the life cycle of a plant depends on its tolerance, supply of the resource and how it is affected by the presence of the second type of annuals. We demonstrate that coexistence of the two species is possible, and we determine the conditions for it.

## 1 Introduction

The problem of coexistence of species and maintaining biodiversity is one of the central issues of plant ecology. As such it has many facets and the number of papers devoted to the subject is enormous. There are many papers discussing the competition - colonisation trade-off (Levins and Culver 1971; Tilman 1994; Holmes and Wilson 1998; Yu and Wilson 2001; Coomes et al. 2002), role of heterogeneity of space (Chesson 2000; Tilman 1987; Pacala 1986) or various type of disturbances, like fire, flooding or grazing (Roxburgh et al. 2004; Miller et al. 2012; Seifan et al. 2012). Pairwise (PW) experiments of annual plants have been reported by Goldberg and Fleetwood (1987) where the effect of the presence of nearest neighbouring plants has been studied and the plant types differed in many aspects. The focus was on the role played by the size of the seeds in the competition between plants. The problem of how the presence of one type of plants affects the target plants, has no satisfactory answer (Grime 1977; Keddy and Shipley 1989; Tilman et al. 1990; Connolly et al. 2001) and therefore further, also theoretical, studies are much needed.

In this paper we present a model of a PW simulation experiment of annual plants. This model, described in detail in the next section is of individual based type and its properties are determined by Monte-Carlo type numerical simulations.

We revisit some important questions (Connolly et al. 2001), namely:

- In a pairwise evolution which species dominates at a particular time?
- Is a coexistence of two species possible?
- What is the effect of another species presence on the performance of the target plants?
- How the above relations change when the supply of the resource vary – from a shortage, through optimal to a surplus. How the dominance changes with the supply of the resource?

Dominance means here that one species is more abundant than the other (Connolly et al. 2001).

There are many factors characterising a plant community and accounting for even most of them would make the model intractable, banning exploration of the full parameter space. Therefore we have decided to reduce the characteristics of the plants to just their demand for one resource, tolerance to a shortage of it and the maximum number of seeds a plants can produce in a year. Once the basic mechanisms of such a system are understood, it would be possible to extend the model by considering different size of the plants, competition for more than one resource, different maximum number of seeds and their size, effect of litter etc, (Kącki and Pękalski 2011; Pękalski and Szwabiński 2013; Droz and Pękalski 2013). Also spatial and/or temporal inhomogeneities either in the form of cyclic or random changes could be incorporated. We study a habitat composed of a population of annual plants living on plaquettes of a square lattice of size *L* × *L* with hard boundary conditions. In the first part of the paper we shall investigate a system with only one type of annual plants. Here we shall determine the role of the control parameters and the mechanisms at play. In the second part we shall study a system with two types of plants and we will establish the influence of the presence of the second type of plants on the first one. In both cases, a lattice site (plaquette) could be either empty or contain only one plant. Initially, a given number of plants is randomly distributed on the lattice. Life cycle of a plant is composed of 15 weeks, making a year. During a year plants are growing and produce seeds according to their fitness, which will then determine the chance a plant has to survive till the moment of dispersing its seeds. At the end of a year seeds are dispersed and the plants die.

External conditions, which could incorporate nutrients, water and light, are summarised in the parameter

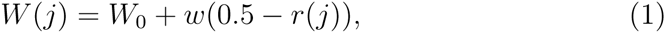

which describes the amount of the resource available during week *j*. To account for possible changes of the resource from one week to another we introduce fluctuations with the amplitude *w* around *W*_0_ and *r*(*j*) is a random number taken from a uniform distribution in the interval [0, 1].

All plants within a given population have the same average characteristics (except their position in the habitat). The features of the plants are:

1. *µ* the maximum number of seeds a plant could produce in one year in optimal conditions.
2. *τ* ∈ [0,1] – plants’ tolerance to the shortage of the resource *W*.
3. *d*, the demand for resources may slightly vary among plants. To account for that we assume that the demand of a plant *i, d*_*i*_, is between *d*−∈ and *d*+∈, where ∈ is the amplitude of the fluctuation. Assuming that there is a single value of the external resource, which is optimal, is certainly an approximation. However, allowing for a range of best conditions would lead to additional parameters and eventually will make the analysis of the results less transparent. The demands are fixed and do not change in time. In the following it will prove useful to operate with *γ* = *W*_0_*/d*, a value of the resource, *W*_0_, normalised by the average demand for it *d*. This way the parameter *γ* is dimensionless.
4. We assume that plants having large tolerance to a shortage of the resource have lesser tolerance to a surplus of it. Therefore for *W* (*j*) *> d*, or equivalently *γ >* 1, the value of *τ* is replaced by (1 − *τ*). It should be noticed that when *τ* = 0.5, the tolerance to a shortage and too much of the resource are the same (see below).
5. If a plant has a nearest neighbour, a part *c* of the resource is blocked by the neighbour. In this way interactions among the plants are accounted for. This may correspond to e.g. blocking by the neighbours’ roots a part of water or to reducing light. Root competition and mechanisms inhibiting access of other roots to resources (called sometimes contest competition or allelopathy) is a well known phenomenon (Schenk 2006). Since there could be a shortage as well as a surplus of the resource the effect of this interactions could be negative or positive (competition or facilitation).

First, we shall show how a population of one type of plants with a given tolerance reacts to changes of the external parameter *γ*. Afterwards we shall determine the influence the second type of plants has on the target ones. The plants of the second second type are almost identical to the first ones, except for one feature - their tolerance to a shortage of the resources.

### 1.1 One type of plants

Our algorithm (Monte Carlo type of simulations) is the following. At the initial time, a certain number (2000) of plants is put randomly on 10^3^ plaquettes of the lattice. Each week plants are randomly selected, their demand for the resource is compared with its availability and the result is a measure of the departure from the optimum. For a plant *i* at week *j* the effective resource available to it has the form

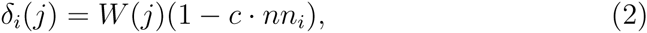

where the factor *c* · *nn*_*i*_ represents that part of the resource which is blocked by *nn*_*i*_ nearest neighbours of the plant *i*. The ratio of the resource available to the plant, *δ*_*i*_(*j*), to the demand for it (*wd*_*i*_), is used in the determination of the plant’s fitness at time *j*

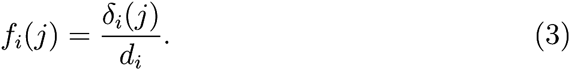

Fitness has a maximum when the supply equals the resource demand for it. When there is too much of the resource, like too much water in the soil, the fitness goes down. In short, fitness peaks in optimal conditions. Fitness from each week, *f*_*i*_(*j*), is averaged over the weeks to yield *f*_*i*_, which is used to calculate the probability of survival of the plant *p*_*i*_ at the end of a year. The form of the probability is obtained from the following empirical considerations:

1. *p*_*i*_ should be 1 if all the needs are met by the resources,
2. *p*_*i*_ should vary smoothly in a way taking into account the tolerance factors.
3. *p*_*i*_ should go to zero when *γ* → 0, because if there is no resources a plant cannot survive.

A simple function fulfilling the above criteria is

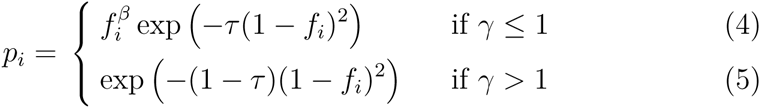

where *β* is a positive number controlling the shape of the probability function for *γ* ≤ 1. If a random number *r*_*i*_ ∈ [0,1], taken from a uniform distribution, is greater than *p*_*i*_, the plant is eliminated. Otherwise the plant survived, and the number of seeds, which it produced, is calculated as a product of the maximum number of seeds *µ* and *p*_*i*_. Using the same formula for checking the plant’s survival and the number of seeds it produced in a year might be a simplification, however, as shown by (Shirley 1929) the two are closely related and therefore such an assumption is well justified. The seeds are randomly distributed on the 13 plaquettes composed by the first, second and third nearest neighbours of the plant *i* and the site *i* itself. After dispersing their seeds plants die.

Next comes springtime when seeds can germinate. Each plaquette containing a seed is randomly selected and the germination test is performed much in the same way as the survival test was run for adult plants. Here however the seeds do not feel the presence of neighbouring seeds, since the seedlings are too small to have an effect on their neighbours. Once all plaquettes with seeds have been visited, the seeds are removed from the system - there is no seed bank. Seedlings become adult plants and a new year starts.

Our choice of several parameters of our model has been motivated by the papers of Hickman (1977) and of Reynolds (1984) who studied annual plants living in mountain areas. The life cycle of such plants is about 15 weeks, they produce up to 10 seeds per plant and the seeds success depends strongly on the external conditions (Reynolds 1984). Plants have also different tolerance to shortage of water (Hickman 1977). Their coexistence has been observed for a period of several years (Hickman 1977). The value of the “interaction strength”, *c* is chosen as 0.1. Larger values could be unrealistic, since it seems that half of the a priori available resource could not be blocked by neighbours. We have verified that making a year last 45 weeks, which would be appropriate for plants living in moderate climates, does not affect the results. As the dynamics is stochastic, some averaging should be performed. Note however that it is also interesting to study the role played by a control parameter (like *τ* or *c*) on the fate of a single population, as will be demonstrated later on.

### 1.2 Two types of plants

A similar approach can be applied in the case where two types of plants, called P1 and P2, are present. The two types of plants could differ in many respects - their demand for the resource, type and number of seeds, tolerance to a shortage of the resource etc. To keep the problem simple we assume that the only difference is the tolerance to a shortage of the resource. The initial conditions are such that a fraction *f*_1_ of the lattice is occupied by the target plants P1 while another fraction *f*_2_ is occupied by plants P2. The fraction *f*_*k*_ (k = 1,2) when multiplied by the initial density determines the density of plants of *k* type. Assuming that initially we put 2000 plants on a lattice with 10000 plaquettes, the fractions *f*_*k*_ vary between 0 and 0.2. Increasing the initial number of plants to, say, 5000 is virtually unnoticed after some 10 years.

We have the following control parameters - reduced resource supply *γ*, tolerance of plants P1 - *τ*_1_ and of plants P2 - *τ*_2_ and finally the initial fractions of two types of plants - *f*_1_, *f*_2_.

The evolution rules for the two types of plants are straight extensions of the ones described above for one type of plants. The only new aspect is concerning seeds germination. After the plants have dispersed their seeds, a plaquette may contain *s*_1_ seeds coming from P1 and *s*_2_ seeds coming from P2. Only one seed will be chosen and submitted to the germination test. This seed is selected according to the majority rule, i.e. the chance of choosing, say, of a P1 seed, is proportional to the fraction *s*_1_*/*(*s*_1_ + *s*_2_) of this type of seeds on the plaquette. It corresponds to the lottery model introduced by Chesson and Warner (1981) with equal weights.

## 2 Results

### 2.1 One type of plants

We consider first a system of dimensions *L*×*L* with *L* = 100 on which initially 2000 plants of one type are put in random positions. Dynamics of this system has been described in the previous section. We have found that a population reaches a stationary state, where the fluctuations of the number of plants are less than 1%, in just several years. The number of plants in the stationary state vary very little from one realisation (random initial distribution of the same number of plants) to another. We have limited the simulations to 100 years and the data presented in this section are averaged over 20 realisations.

Figure 1 shows the number of plants as a function of *γ* = *W*_0_*/d*, the resource supply *W*_0_ divided by the average demand for it *d*, for several values of the tolerance *τ*. As could be expected, for small tolerance, like *τ* = 0.3, plants need larger values of *γ* to survive than when the plants tolerance is greater, but on the other hand in this case they can survive when *γ* is much larger than 1. Maximum of the number of plants does not depend on the tolerance and is located at *γ* ≈ 1.2. The shift from *γ* = 1 is due to the fact that part of the resource is blocked by the neighbours of each plant, hence *γ* should be larger than 1.

**Figure 1:**
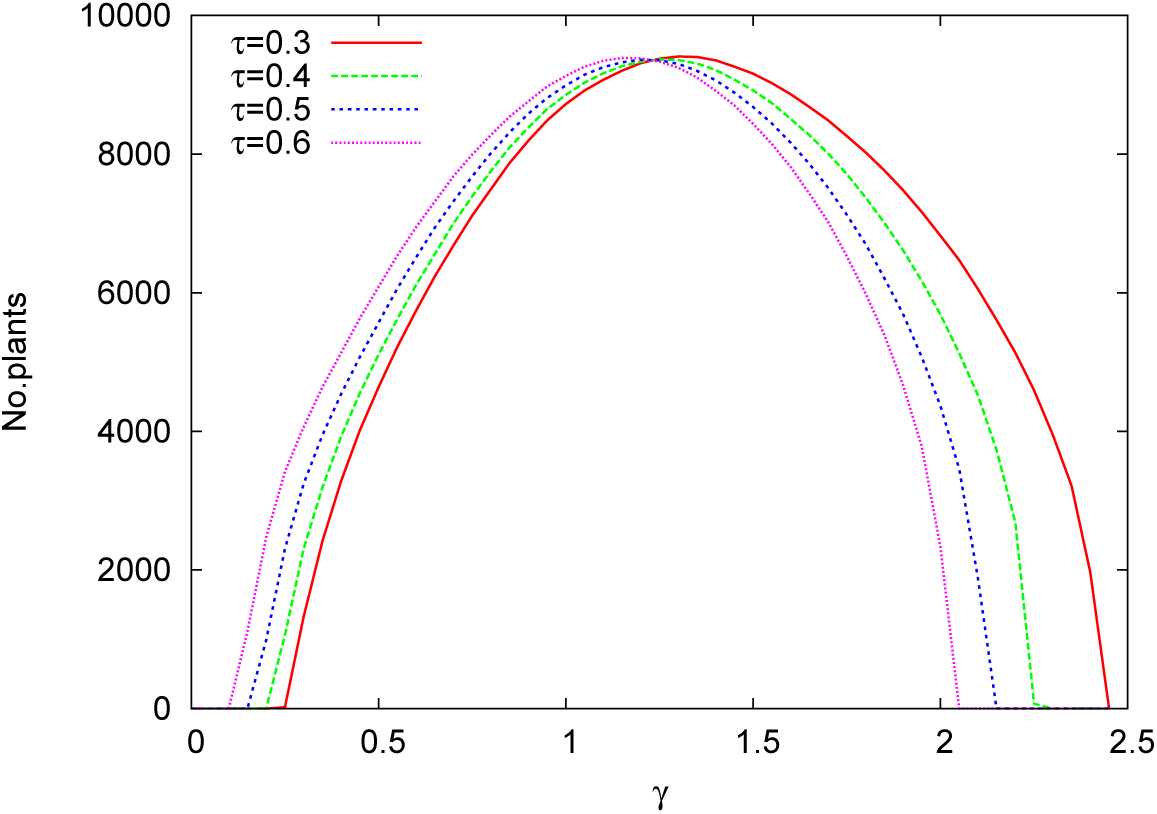
Number of plants P1 for various tolerances as a function of the reduced resource supply *γ*. Colour on line.

Next, figure 2 shows the three successes - *S*_1_, *S*_2_, *S*_3_ defined as follows. *S*_1_ is the seeds success, i.e. the ratio of the number of seedlings in a given year to the number of plaquettes with seeds in the same year. Since from a plaquette only one seed could germinate, it is the number of plaquettes, not the number of seeds, which determines this success. *S*_2_ is the seedlings success - the ratio of the number of adult plants which were able to produce seeds, to the number of seedlings. Finally, *S*_3_ is the total success - the ratio of adult plants producing seeds to the number of plaquettes with seeds, or simply *S*_3_ = *S*_1_ · *S*_2_. We have found that the three successes depend only very weakly on the value of the tolerance, hence only the plot for *τ* = 0.5 is presented. Which success dominates and which determines the overall one, depends on the value of *γ*, i.e. the supply of the resource.

**Figure 2:**
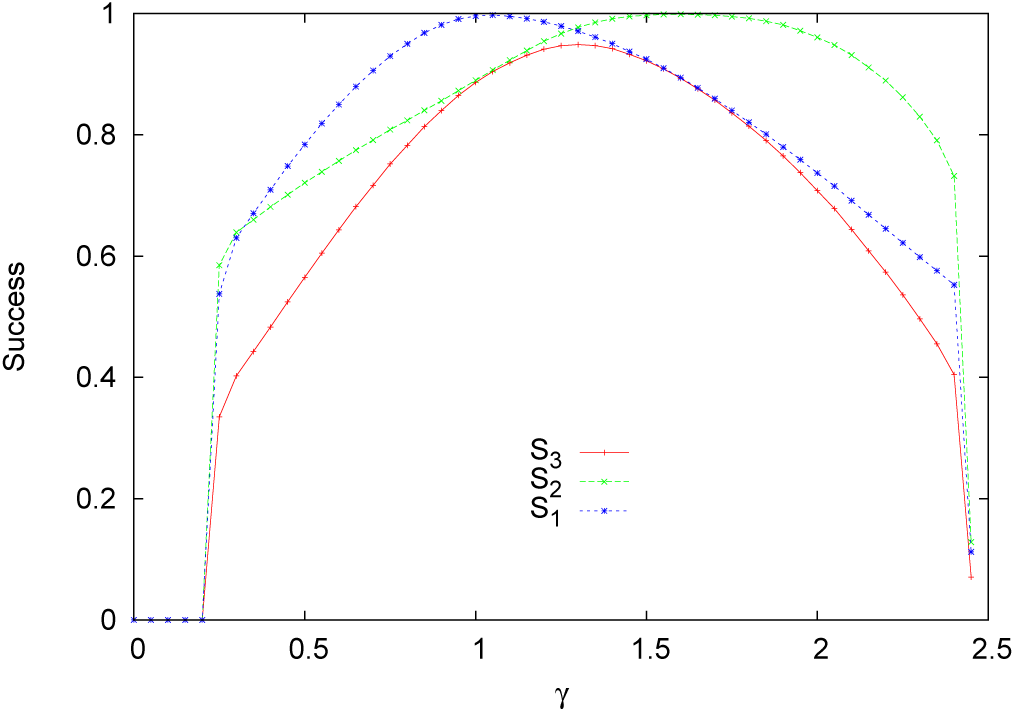
Three successes for P1 plants. *S*_1_ - seeds success, *S*_2_ - seedlings success, *S*_3_ - total success of plants (product of the previous ones) as functions of the resource supply *γ* and for *τ* = 0.5. Colour on line.

Spatial arrangement of plants may change in different external conditions, viz. for different *γ*. A convenient way to measure it is the ratio of the average number of nearest neighbours 〈*NN*〉 to the average number of next nearest neighbours 〈*NNN*〉. When there is not enough resource (low *γ*) plants are expected to avoid growing in closely packed clusters, since each NN blocks a certain amount of, already scarce, resource. When the supply of the resource is about what a plant needs, one may expect a balance between the average number of 〈*NN*〉 and 〈*NNN*〉. Finally, when there is too much of the resource, growing in close clusters becomes beneficial, as it allows to reduce the too large supply. This is exactly what is seen in figure 3.

**Figure 3:**
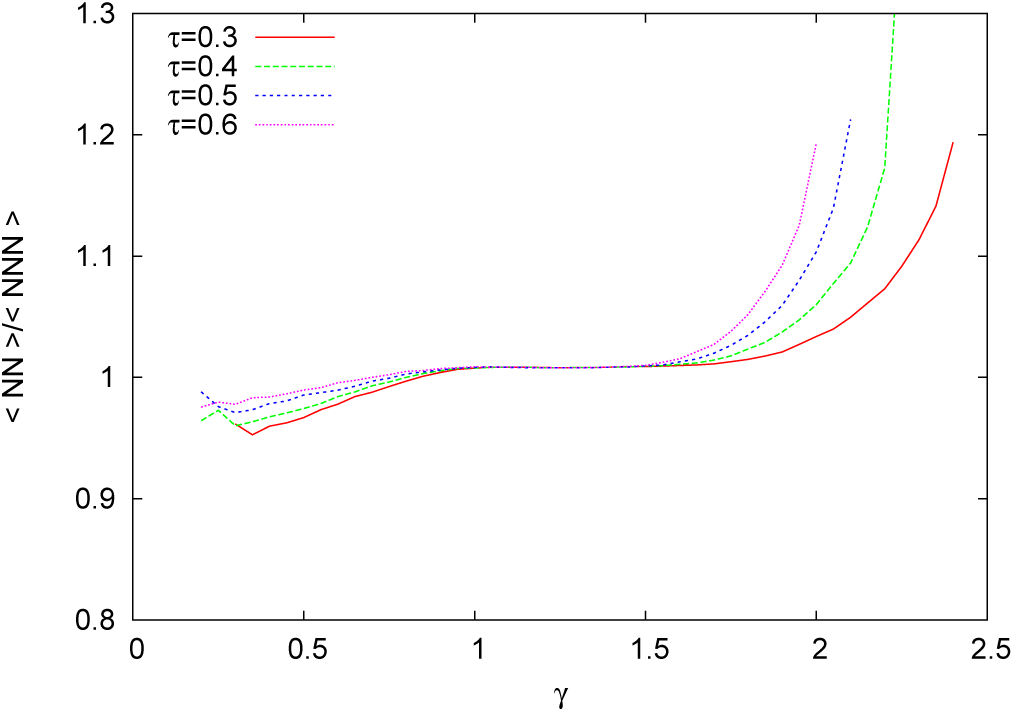
〈*NN*〉*/*〈*NNN*〉 for several tolerances *τ* as a function of the resource supply *γ*. Colour on line.

The presented above plots allow the following description of the basic mechanisms of our model. There is a tendency of plants of adjusting their spatial organisation according to the external conditions in order to cope with either surplus or shortage of the resource. This in turn determines whether the elimination will take place at the seedlings or adult plants stage. If *γ <*1, hence there is a shortage of the resource, eliminated are mostly adult plants. It could be explained as follows. When the resource is scarce, seedlings, which do not have their access to the resource blocked by their NN, are better off than adult plants which loose a part of the resource supply. This lost may be crucial and an adult plant could be eliminated before producing seeds. When *γ >* 1, there is too much of the resource and the elimination is at the level of seeds. Once a seed germinated and developed into an adult plant, its conditions are better since part of the surplus of the resource is blocked. The total success of plants is therefore determined mainly by the success of the adults plants when *γ <* 1 and by the success of seeds when *γ >* 1.

## Two types of plants

After showing the basic mechanisms of our model, we may turn into applying it to a more complex case, namely two, called P1 and P2, quite similar types of annuals living in the same habitat. We assume that both types are present there from the beginning of the study and no immigrants are allowed.

We assume that the initial fractions of the P1 and P2 are equal. Such an assumption allows us to estimate whether a given species contributes more at the end than at the beginning and then we can tell which of the species outperformed the other (Connolly et al. 2001). It corresponds to the average relative growth rate concept (Evans 1972). We shall call the P1 plants the target plants with fixed value of the tolerance at *τ*_1_ = 0.5 and we shall investigate how the presence of P2 of various tolerances, influences the existence of P1. Fixing the value of *τ*_1_ and changing only *τ*_2_ is reasonable, since we know from the previous section how the dynamics of P1, living alone, changes with *τ*_1_ and moreover the only difference between P1 and P2 are their tolerances. When two types of plants are present a stationary state is also attained, although it corresponds to just one type of plants alive. The less adapted species are sooner or later eliminated. This may happen after a long time, like 800 years, or just after 50 years and the outcome depends on the supply of the resource and initial arrangement. Elimination of a worse adapted species has been established before (Levin 1970; Chesson and Huntly 1997; Chesson 2000). It, is of course quite unrealistic to assume that external conditions will remain the same for 800 years. Therefore we have decided to study the case of two types of plants during an interval of 100 years. Clearly at that time the system is not in a stationary state but in a transient one. Such a period is still quite long from an experimental point of view. How the picture changes when another time limit is chosen, will be discussed later. The figures for two types of plants for the three successes and the dependence of the average number of NN divided by the average number of NNN are quite similar to the Figures 2 and 3, and therefore are not presented.

For one type of plants averaging over just 20 realisations gave smooth curves. Dynamics of a system of two types of plants is much more complex and we had to average over 200 realisations to arrive at the figures presented in the next section.

From figure 4 we see that the presence of P2 has a profound effect on P1, restricting their existence range and changing the abundance. There are regions when the better adapted plant eliminates the other one, but there is also an interval of the values of *γ* where coexistence is possible. The region is larger when the tolerances of P1 and P2 are more similar. It should be noted that even when the tolerances are very close, like *τ*_1_ = 0.5, *τ*_2_ = 0.45, the worse adapted species is eliminated in a large interval of the values of the resource which were acceptable when there was just one type of plants. This effect is better seen in figure 5 where we compare the number of plants P1 when they grow without P2 (the *mono* curve) and when P2 are present and have various tolerances. Several features should be noticed. When *τ*_2_ = 0.3 the P1 are unaffected at all for small values of *γ*, since there the advantage of P1 is so large that P2 are eliminated. Once however the P2 are able to survive, they reduce the number of P1 and with growing *γ* the process of replacing P1 by P2 is more and more fast. If *τ*_2_ = 0.45, the P2 appear at smaller values of *γ* than when *τ*_2_ = 0.3, but could eliminate P1 only when *γ* is quite large.

**Figure 4:**
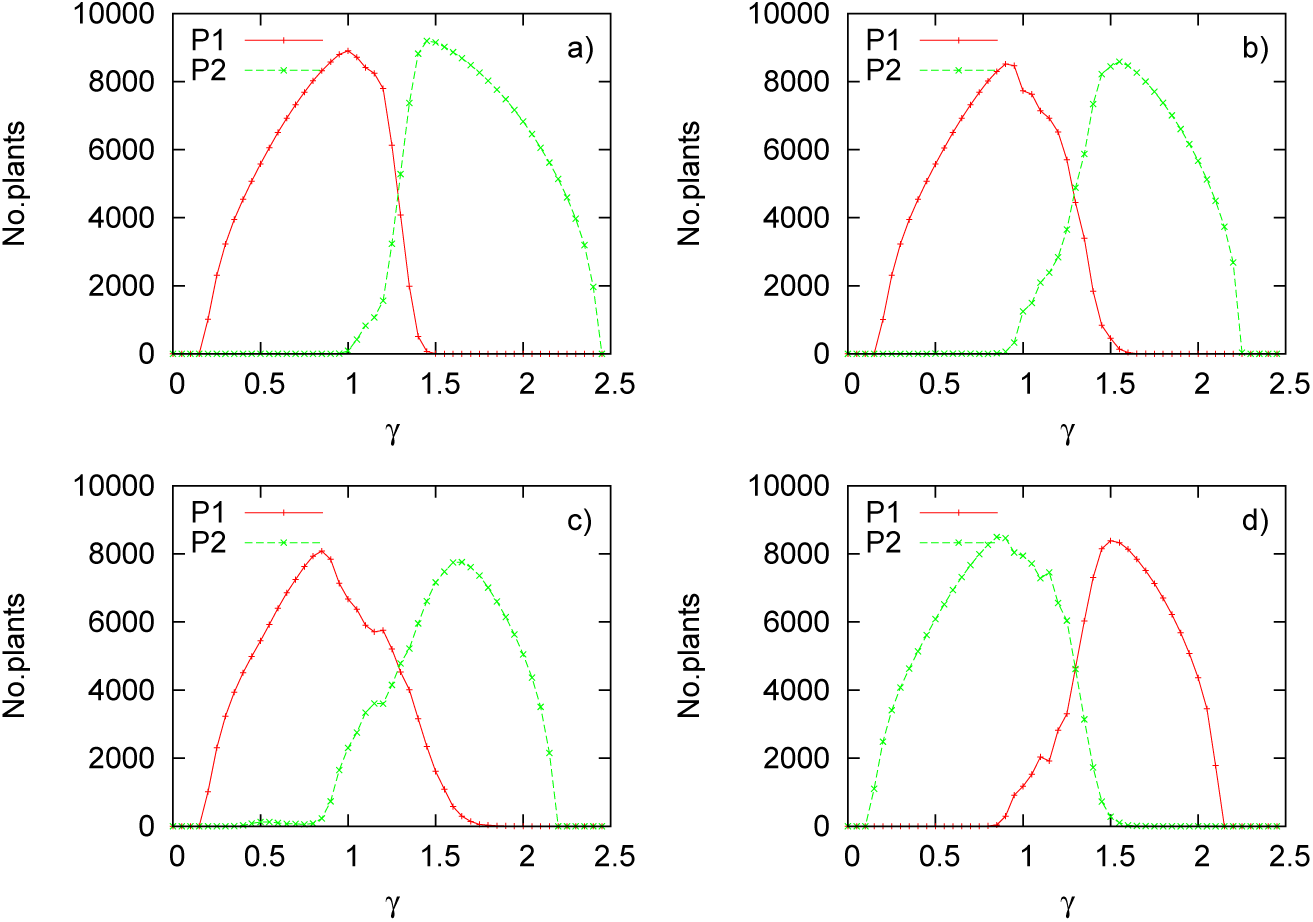
Number of P1 and P2 plants for various tolerances of P2 and as a function of the resource supply *γ*. *τ*_1_ is everywhere equal 0.5. a) *τ*_2_ = 0.3, b) *τ*_2_ = 0.4, c) *τ*_2_ = 0.45, d) *τ*_2_ = 0.6. Colour on line.

**Figure 5:**
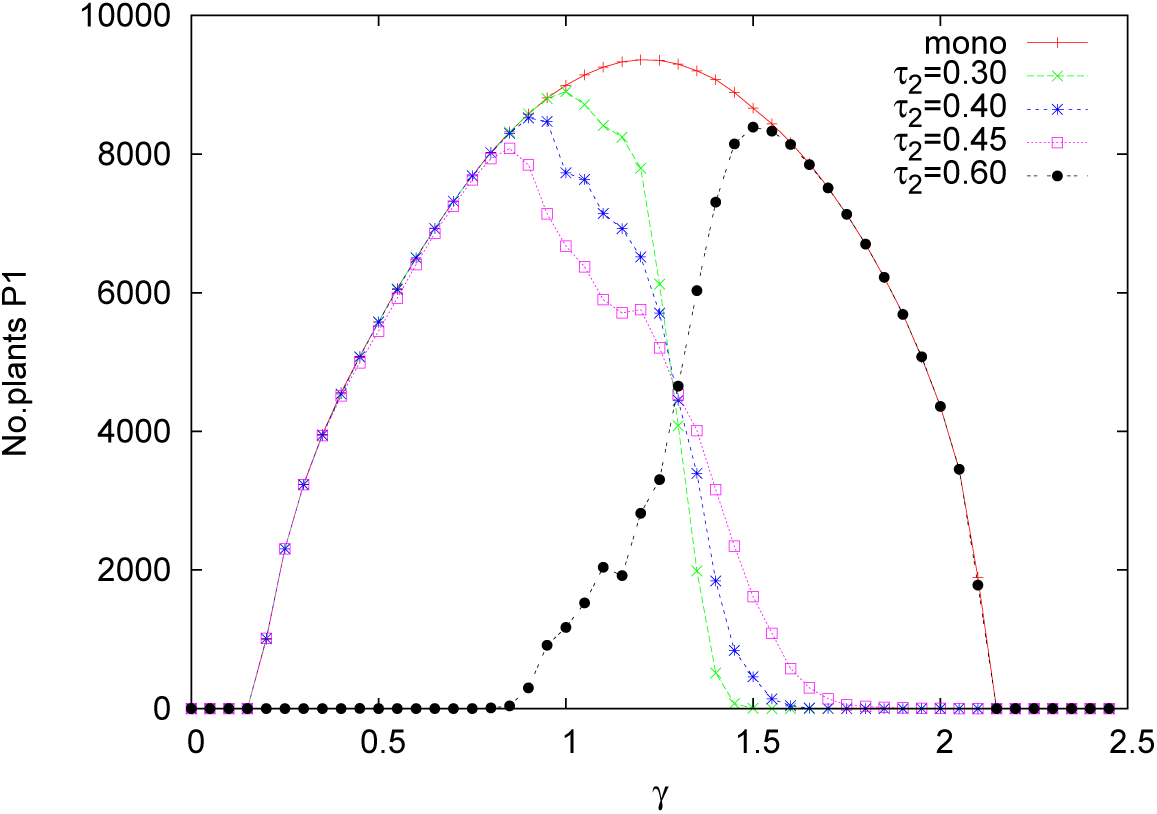
Number of P1 where they grow alone (*mono* curve) and with P2 of various tolerances as a function of the resource supply *γ*. *τ*_1_ = 0.5 for all cases. Colour on line.

This figure gives an answer to the question often asked by ecologists (Connolly et al. 2001) - how much the target species performance (here the number of plants) is attributable to other species presence. The figure shows how this influence changes with the supply of the resource and the relation between tolerances of the two species.

When two species coexist and compete for a limited resource, often a plant competition index is introduced. There is however still a controversy how it should be defined. In a paper by Weigelt and Jolliffe (2003) nearly 50 of them have been described and discussed. In view of the controversy we have chosen a simple one, namely the Relative Competition Index (RCI), introduced by Grace (1995) as

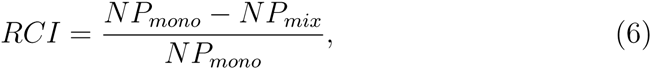

where *NP*_*mono*_ is the number of plants in a monoculture and *NP*_*mix*_ is the number of the same type of plants living with other plants. In Figure 6 the RCI is presented as a function of the resource supply *γ* and several values of *τ*_2_. There is no competition when RCI is either zero (P2 plants are there eliminated) or when RCI = 1, when the P1 plants cease to exist. The region in between marks the competition and coexistence range. If RCI ≤ 0.5 the P1 are dominant, otherwise P2 are more abundant.

**Figure 6:**
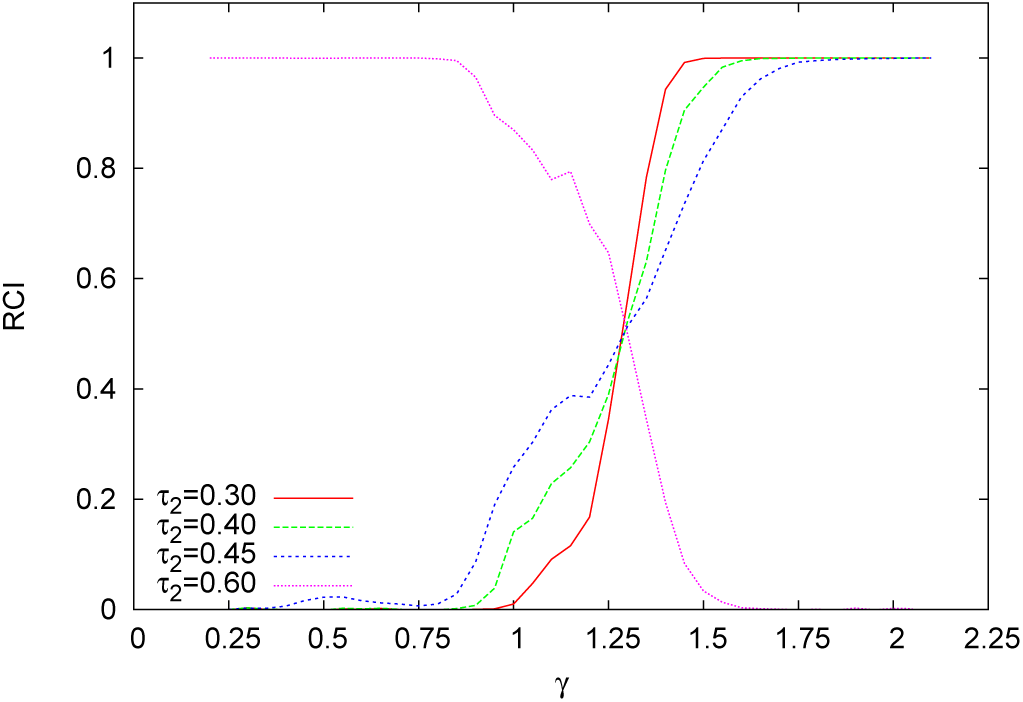
The RCI index, for definition see the text, as a function of the resource supply *γ* and several values of the P2 plants tolerance. Colour on line.

An interesting problem, which could be studied only via computer simulations in agent based models, is the spatial organisation of the two populations. Figure 7 shows it after 100 years for *γ* = 1.4, *τ*_1_ = 0.5, *τ*_2_ =0.4, both in a whole lattice and in a blow-up showing more details of the spatial structure. One sees clearly that plants of the same type have a tendency to grow in clusters. There are several ways to measure this effect. One of them is presented in Figure 8 as the average number of NN of the same type as the central plant (*same* line) and of both types (*both* line). It is evident that e.g. for plot a), when *γ* is very small, the P1 plants have only NN of the same type, as there are no other plants in the system. However for *γ* ≈ 1.3 there is already a large number of P2, yet still the P1 have mostly P1 as their NN.

**Figure 7:**
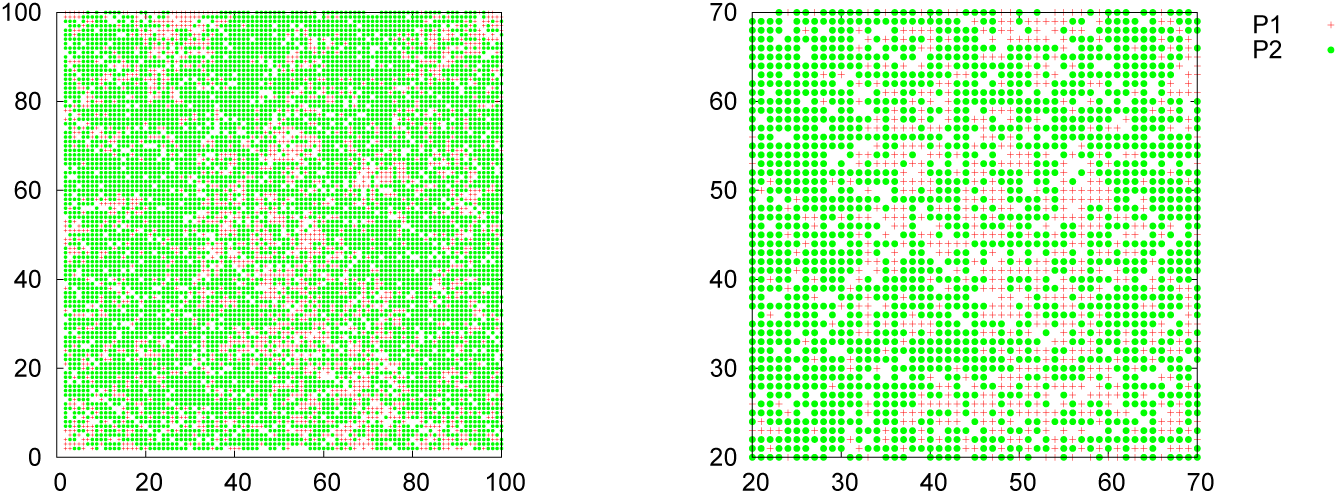
Spatial arrangement of the P1 (red crosses) and P2 (green circles) plants at the end of simulation (100 years) for *γ* = 1.4, *τ*_1_ = 0.5 and *τ*_2_ = 0.4. Full lattice (left) and a part of it (right) Colour on line.

**Figure 8:**
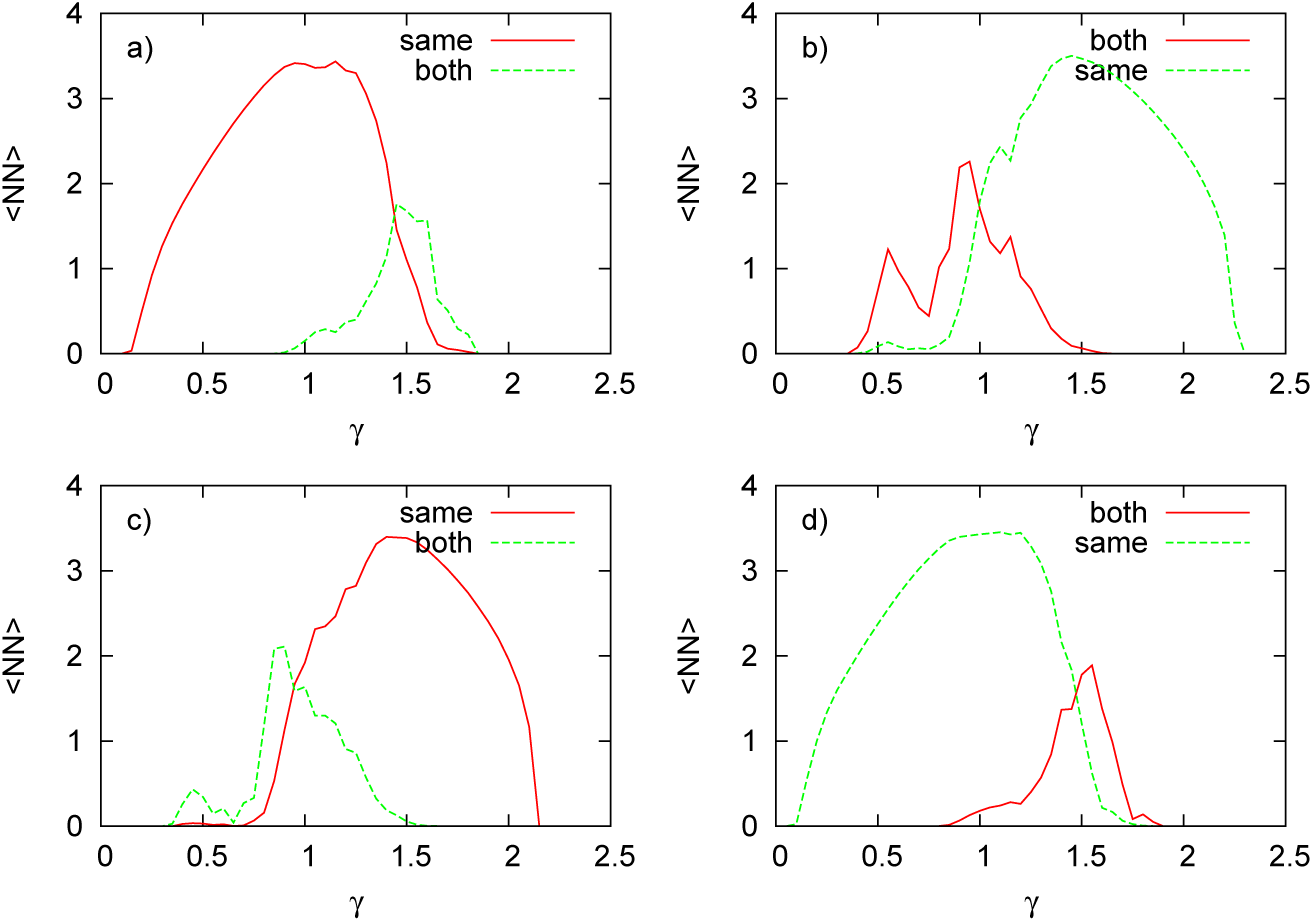
Average number of nearest neighbours, as functions of the resource supply *γ*, of the same type (*same*) and of both (*both*) types for P1 and P2 and two values of *τ*_2_. a) P1 plants *τ*_2_ = 0.4, b) P2 plants *τ*_2_ = 0.4, c) P1 plants *τ*_2_ = 0.6, d) P2 plants *τ*_2_ = 0.6. Colour on line.

Another way to see possible regularities in the spatial arrangement of plants, is to plot the number of plants of a given type along some direction. Figure 9 shows it for the X-axis, *γ* = 1.4 and *τ*_2_ = 0.4. The total number of plants along that axis is, *grosso modo*, constant, P2 being more numerous, but we see also that in some areas the number of P1 grows at the expanse of P2, which indicates the presence of a P1 cluster.

**Figure 9:**
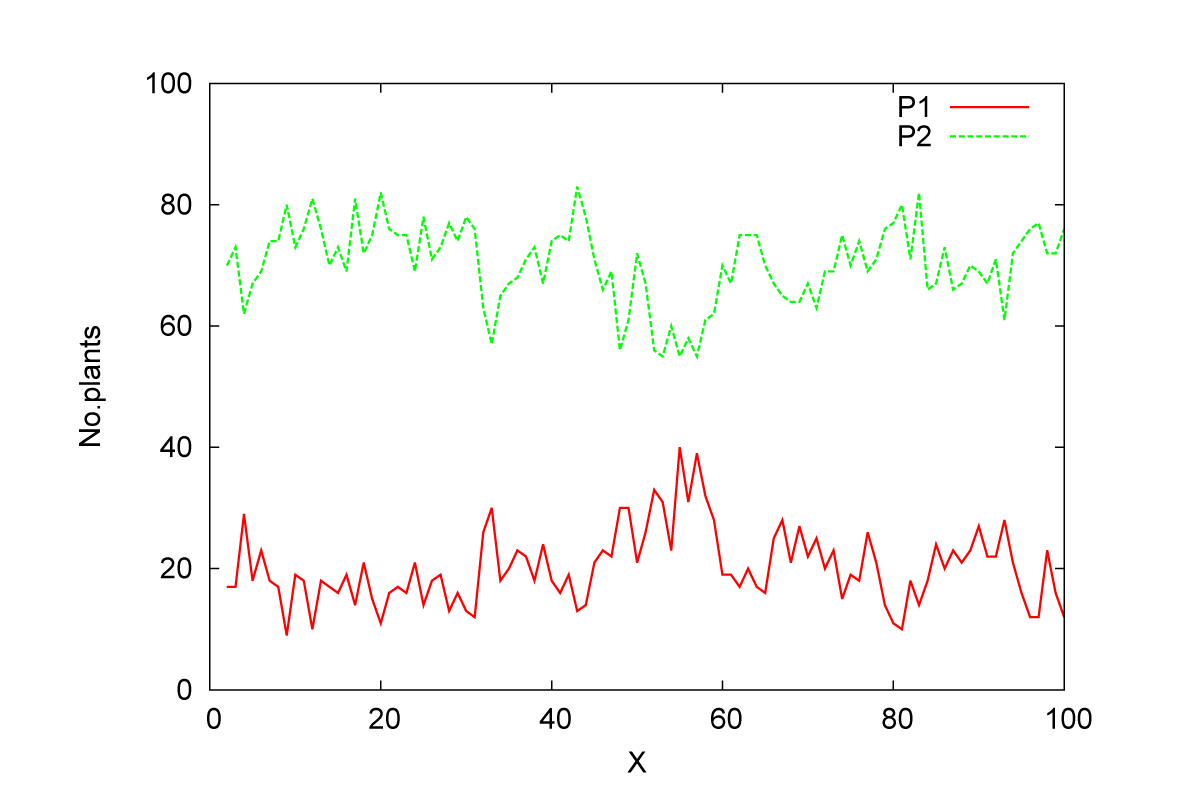
Number of P1 and P2 plants along the X-axis at *t* = 100 years and for *γ* = 1.4, *τ*_1_ = 0.5 and *τ*_2_ = 0.4. Colour on line.

When the tolerances of both types of plants are similar, like *τ*_1_ = 0.5, *τ*_2_ = 0.45, the number of both types of plants are comparable and the effects of appearance of local clusters are stronger (see Figures 10 and 11). Comparison of the two Figures 9 and 11 shows also that when the tolerances *τ*_1_ and *τ*_2_ are getting closer, the clusters of the same type of plants are growing.

**Figure 10:**
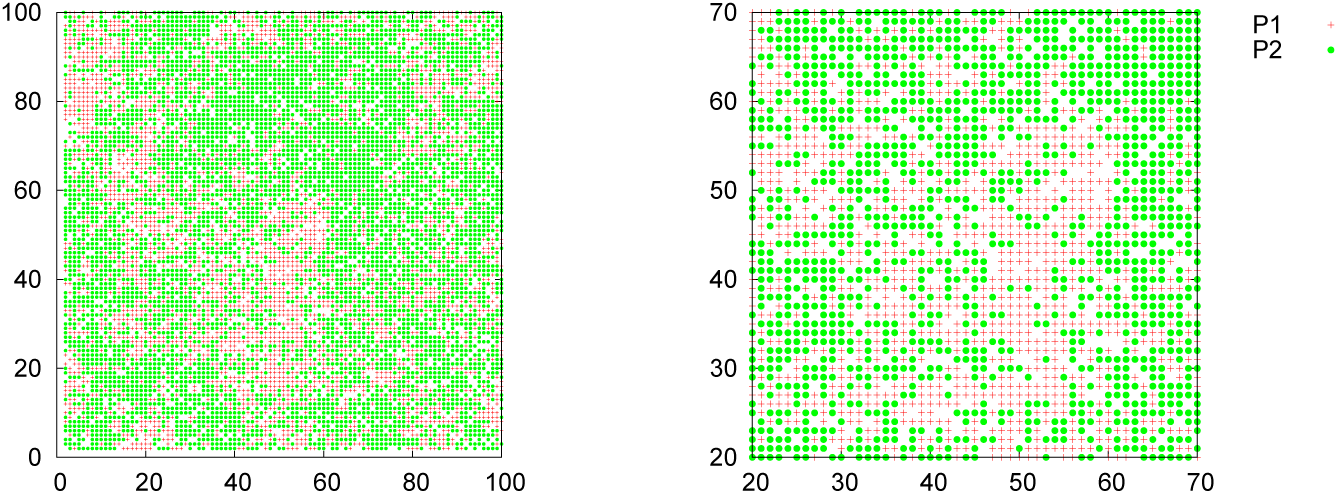
Spatial arrangement of the P1 (red crosses) and P2 (green circles) plants at the end of simulation (100 years) for *γ* = 1.4, *τ*_1_ = 0.5 and *τ*_2_ = 0.45. Full lattice (left) and a part of it (right). Colour on line.

**Figure 11:**
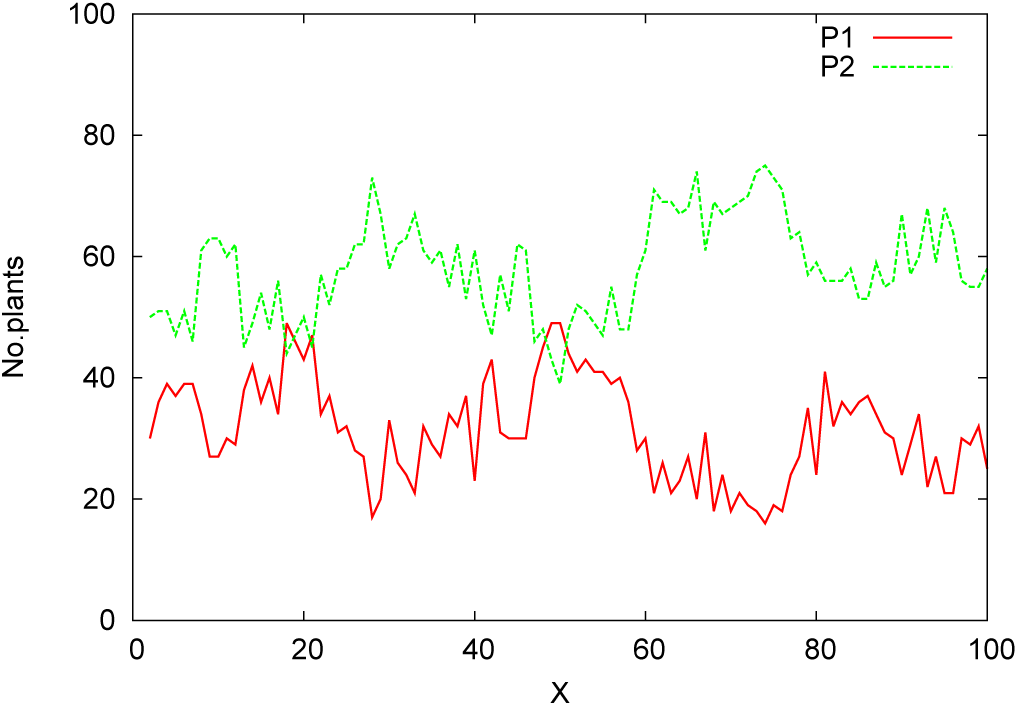
Number of P1 and P2 plants along the X-axis at *t* = 100 years and for *γ* = 1.4, for *τ*_2_ = 0.45. Colour on line.

Finally, we discuss how the results presented above change when the parameters we have fixed, take another values. Since the plots for the two types of plants were obtained for a transient regime, it is important to check how the picture will change would we take another time limit at which the plants’ abundances were calculated. 100 years taken by us in the simulation, is a long period and therefore we have repeated the simulations for 50 years time limit. There are some, but quite minor, differences between the results for the two time limits. We may therefore conclude that in the transitory regime at which we investigate the population evolution, the results do not change qualitatively when the time limit varies within, say, the interval [30 - 150] years and in this respect the results are generic. The scatter of the reduced demand, ∈ = 0.2, for the resource, makes the maximum number of plants as a function of *γ* less sharp, however a difference between ∈ = 0 and ∈ = 0.2 is rather small. Likewise the assumption that the supply of the resource fluctuates from week to week with an relative amplitude *w/W*_0_ = 0.02, has a small effect and its role is rather elimination of unrealistic repeating each week the same supply of the resource. Similarly, taking ∈ ≠ 0 prevents us from considering a system of clonal plants.

The size of the system *L* = 100 is a compromise between execution speed and avoiding stochastic extinctions and poor statistics. For small systems, like *L* = 20, even local demographic stochasticity could, in the case of non-linear dynamics, lead to global effects (Bolker and Pacala 1999; Chesson 2000; Durrett and Levin 1994).

There are two other parameters which could influence the dynamics of plants in our model. First is *c* - the “interaction strength” describing the effect on a plant of its NN. When *c* = 0 we have a system of completely independent plants when the neighbourhood have no influence on the central plant. The main difference among plots for various values of *c* is the shift of the maximum number of plants towards larger values of *γ* when *c* is growing, since with increasing *c* more and more of the resource is blocked by the NN. The second effect of growing *c* is pushing down the maximum abundance, at *γ*_*max*_, and at the same time increasing the abundance for *γ* close to *γ*_*max*_. This comes from the fact that for large *c* the nearest neighbourhood plays more important role and since not all the plants have the same neighbourhood, optimal conditions are more widely spread than for small values of *c*. Also the coexistence region shrinks with growing *c*. For *c* = 0 the dependence of the number of plants on *γ* has a very sharp maximum at *γ* = 1.

The last parameter *β >* 0 is needed to assure that the probability of survival is vanishing when there is no resources. We have verified that changing *β* from 0.1 to 0.5 in our simulations does not influence the results in a significant way. Small values of *β* produce a smoother probability distribution for *γ* ≤ 1.

## 3 Conclusions

We have presented and discussed a computer based simulation model describing life cycles of a population of annual plants competing for a single resource. Two scenarios have been investigated. In the first one only one type of plants lived, while in the second there were two types. Plant characteristics have been reduced to their tolerance to a shortage or surplus of the resource and the maximum number of seeds a plant could produce in one year. The only difference between the two types of plants is their tolerance to the shortage of the resource.

The questions we have asked are – is a coexistence of two different species possible in such a simple system, without the competition-colonisation trade-off (Coomes et al. 2002; Yu and Wilson 2001) or disturbances (Chesson 2000; Miller et al. 2012; Seifan et al. 2012). If so, how the dominance of one species depends on the supply of the resource and the difference in the tolerances. How the presence of a second type of plants, with quite similar characteristics, affects the target plants. What will be the spatial structure in the coexisting phase and how it will depend on the control parameters. To the best of our knowledge such investigations have not yet been carried out.

The results obtained from the simulations and presented in the Results section, allow us to conclude that our simple model leads to a coherent description of the situation and bring new light on the basic questions listed above.

While the system of one type of plants arrives at a stationary state very fast (up to 10 years), dynamics of a mixture is by far more complex. The final state, which could be attained sometimes only after several hundred years, is always a single species state. In the transient period coexistence, lasting also sometimes very long, is possible. The range of values of the resource over which it happens, is larger if the tolerances do not differ much and happens where the supply is close to the demand. Growth rates of the population depend very strongly on the supply of the resource and the difference of the tolerances. Where the two are close, the growth rate of the target plants is more affected by the presence of the second type of plants than when the tolerances differ more. Life cycle of a plant could be controlled either by elimination of the seedlings (including germination blocking) or by elimination of adults plants. We have shown that it depends on the supply of the resource which of the mechanism is more important. The value of the tolerance does not play a significant role here.

Using individual-based model allowed us to determine also spatial organisation of plants. We have shown that depending on the availability of the resource the plants have a tendency to stay away one from another when the resource is scarce. When there is an adequate amount of it, there is no visible pattern and when there is too much of the resource, the plants have a tendency to form dense clusters. In a system of two types of plants they form groups of alike plants and this tendency is growing when the difference in the two tolerances is getting smaller.

